# Rapid, raw-read reference and identification (R4IDs): A flexible platform for rapid generic species ID using long-read sequencing technology

**DOI:** 10.1101/281048

**Authors:** Joe Parker, Andrew Helmstetter, James Crowe, John Iacona, Dion Devey, Alexander S. T. Papadopulos

## Abstract

The versatility of the current DNA sequencing platforms and the development of portable, nanopore sequencers means that it has never been easier to collect genetic data for unknown sample ID. DNA barcoding and meta-barcoding have become increasingly popular and barcode databases continue to grow at an impressive rate. However, the number of canonical genome assemblies (reference or draft) that are publically available is relatively tiny, hindering the more widespread use of genome scale DNA sequencing technology for accurate species identification and discovery. Here, we show that rapid raw-read reference datasets, or R4IDs for short, generated in a matter of hours on the Oxford Nanopore MinION, can bridge this gap and accelerate the generation of useable reference sequence data. By exploiting the long read length of this technology, shotgun genomic sequencing of a small portion of an organism’s genome can act as a suitable reference database despite the low sequencing coverage. These R4IDs can then be used for accurate species identification with minimal amounts of re-sequencing effort (1000s of reads). We demonstrated the capabilities of this approach with six vascular plant species for which we created R4IDs in the laboratory and then re-sequenced, live at the Kew Science Festival 2016. We further validated our method using simulations to determine the broader applicability of the approach. Our data analysis pipeline has been made available as a Dockerised workflow for simple, scalable deployment for a range of uses.

## Introduction

DNA-based taxonomic identification has become widespread since the characterisation of “universal”, easily amplifiable and sufficiently variable DNA barcoding genes, such as COI, rbcL and matK (Hebert *et al.* 2003, 2016a; CBOL Plant Working Group *et al.* 2009; Coissac *et al.* 2016). In many circumstances the level of taxonomic resolution that short DNA barcodes can provide is suitable for the chosen purpose, but it has become clear that barcoding genes cannot always provide truly species-level discrimination and identification(Hebert *et al.* 2003; CBOL Plant Working Group *et al.* 2009; Little 2011; Hawlitschek *et al.* 2016). Barcoding approaches have certainly been effective for generic level identification, but resolution at the species level is less consistent (CBOL Plant Working Group *et al.* 2009; Little 2011; van Velzen *et al.* 2012; Collins & Cruickshank 2013; Mallo & Posada 2016); Pryron, 2015); and it is evident that assuming that unique barcodes or bins are synonymous with distinct species is potentially misleading (Tang *et al.* 2012; van Velzen *et al.* 2012; Collins & Cruickshank 2013; Schmidt *et al.* 2015; Hawlitschek *et al.* 2016). Hebert *et al.* (2016b) conducted extensive barcoding of the particularly well studied insect fauna of Canada (33 572 species) sequencing of more than 1 million samples (14% of samples produced no barcode) assigning barcodes to family or genus level. Assuming that each unique barcode represented a different species, species richness was estimated at three times the taxonomically described number. This figure may be accurate, but the discrepancy highlights that there number of reasons to believe that barcoding approaches are susceptible to errors introduced by low inter- and high intra-specific genetic variation in different groups (Tang *et al.* 2012; van Velzen *et al.* 2012; Mallo & Posada 2016; Liu *et al.* 2017). Broader, genome-scale identification methods hold a great deal of potential to bridge this gap. These approaches remove ambiguity from species-level identifications and produce information rich data that may be more applicable to species delimitation (Coissac *et al.* 2016; Hollingsworth *et al.* 2016; Parker *et al.* 2017). Indeed, we have previously shown that field-based, genome scale sequencing is able to provide accurate species identification in closely related *Arabidopsis* species (Parker *et al.* 2017). This study used carefully curated whole genome sequences of *A. thaliana* and *A. lyrata* as a reference database (against which newly generated long-read data generated in the field was matched) to explore the potential of this type of data for species identification.

All types of accurate taxonomic identification from DNA sequence data are reliant on reference databases of DNA sequences obtained from specimens identified morphologically by experts. This is exemplified by the rigor with which the most high quality DNA barcode databases are constructed (IBOL)(Hebert *et al.* 2016a) and highlights the important role that taxonomist, taxon-specific expertise and collections based institutions (such as natural history museums and botanic gardens) have played and will continue to play in expanding the utility of DNA-based identification. Increasingly high throughput methods of producing reference DNA barcodes are being developed, but the rate at which reference genomic data is published is substantially slower. Currently, whole genome sequences are available in public repositories (INSDC) for only 2733 eukaryotic species and these are distributed unevenly across the tree of life – 874 bilatarians, 274 plants and 1237 fungi. This means that there are not enough sequences to represent each family of organism, let alone genus and species. As a result, the use of genome scale data for accurate species identification as in Parker (2017) is apparently a very long way away from having the same applicability as DNA barcodes.

Portable single molecule, real-time DNA sequencers have now become a commercial reality. Their potential ubiquity, coupled with cloud computation, mean that bioscience is set to undergo a transformation: millions of researchers, clinicians, conservation professionals and citizen-scientists will have the potential to sequence and analyse genomic material anytime, anywhere (Erlich, 2015). Portable sequencers such as the MinION allow DNA-sequencing to happen anywhere, in real-time, with important applications in human, animal and plant health; border control; resource management; and environmental monitoring. Demonstrated applications have included epidemic monitoring in Guinea (Quick *et al.,* 2016), extremophile sequencing in Antarctica (Michael *et al.*, 2017), assembly of complete plant genomes (Johnson *et al.*, 2017), species ID and phylogenomics in the field (Parker *et al.*, 2017). Sequencing reads from these platforms are orders-of-magnitude longer than those generated by more established, sequencing-by-synthesis technologies. Few tools or techniques are currently optimised for this technology and these developments present opportunities to revisit existing orthodoxies, analyses and uses of DNA data.

In this era of mobile genomic sequencing, can genome-scale reference datasets be produced at a rate that species-level ID from genome resequencing will become a practicable and widely applicable reality? High quality reference genomes are still challenging and time consuming to produce, despite the incredible pace at which new data can be generated. However, the results of our earlier work (Parker *et al*., 2017) suggest that fully annotated and carefully analysed whole genome sequences may not be necessary for species ID from shotgun long read resequencing data. Simulations in which the *Arabidopsis* genomes were fragmented in silico demonstrated that even quite poor quality genome assemblies would be sufficiently information rich for confident identification. This raises the possibility that lab or field generated, genome-shotgun long read data itself may be able to act as a sufficiently detailed sketch of the genome for future identification, which we term rapid-raw-read-reference for ID, or ‘R4IDs‘. In this case, re-sequencing of samples from the same species would still capture enough overlapping sequence with the genome sketch that species identification would be unambiguous. This opens the possibility for rapid and simple generation of a genomic reference database to match the scale of barcode databases, with the added benefit that reference sequences can be generated in the field by expert taxonomists, conceivably at the points at which new species discoveries are made.

Our approach to live identification by real-time DNA sequencing and analysis comprises three steps: a ‘*R4ID*-*training‘* step, in which samples of known origin are sequenced as low coverage whole-genome shotgun libraries and converted to a panel of BLAST databases (being optionally combined with public assemblies); a ‘*R4ID-resequencing‘* step, in which samples of unknown origin are resequenced, also as whole-genome shotgun libraries, and aligned in real-time to the R4IDs databases using BLASTN; and a *‘R4ID-visualisation‘* step by which live results are displayed as a web application with a graphical user interface (GUI). The aim of the R4IDs sequencing in the training step is to provide low-coverage data for accurate ID, not complete genomic sequencing for assembly (although this long-read data could be used to augment existing datasets for this purpose). The aim of the present study was to determine how much R4ID (training) and resequencing (test) data needs to be collected for an accurate ID in reasonable time, both empirically and by simulation.

## Methods and results

### Sequencing and live ID

In the week prior to the Kew Science Festival in 2016, we extracted genomic DNA used Qiagen DNEasy Plant Miniprep kits from five vascular plant species: *Sorbus aria* (Rosaceae); *Beta patula* (Amaranthaeae); *Nepenthes alata* (Nepenthaceae); *Erycina echinata* (Orchidaceae); and *Silene uniflora* (Caryophyllaceae). Sample details, ploidy level and estimated genome size (taken from the Kew C-values Database) are given in Table 1. Estimated genome sizes for these species, or congenerics, range from 0.28pg to 1.9pg.

**Table 1:** Rapid-raw-read-reference identification (R4ID) sampling details. Estimated genome sizes for diploid genomes given in pg, derived from flow cytometry in the Kew C-Values Database (http://kew.org). The closest available species is given where no exact species match was available. *Taxonomy unclear: synonym: *B. vulgaris ssp patula.* (Aiton 1789).

| Clade | Family | Genus | Species | Common name | Tissue | Source | Estimated genome size, pg |
| --- | --- | --- | --- | --- | --- | --- | --- |
| Eudicots | Nepenthaceae | *Nepenthes* | *alata* | Pitcher plant | Leaf / pitcher | Living collection accession (2003-2475) | 0.28 (*N.pervillei*) |
| Eudicots | Rosaceae | *Sorbus* | *aria* | Whitebeam | Leaf | M. Fay (pers. comm.) | 0.71 |
| Eudicots | Caryophyllaceae | *Silene* | *uniflora* | Sea campion | Leaf | ASTP collection | 2.7 (*S.latifolia*) |
| Eudicots | Amaranthaceae | *Beta* | *patula* | Wild beet* | Leaf and stem | M. Chester (pers. comm.) | 1.25 (*B.vulgaris*);<br>1.28 (*B.maritima*) |
| Monocots | Orchidaceae | *Erycina* | *echinata* | The Hedgehog Erycina | U/K | RBG Kew DNA Bank | 1.9 (*E.diaphena*) |

To prepare rapid, raw-read reference ID datasets (’R4IDs‘), whole-genome shotgun libraries were prepared for R9 Oxford Nanopore MinION sequencing using rapid-family (SQK-RAD001) transposase-based kits and protocols, which require less than 20 minutes to prepare. Samples were sequenced for variable run times and basecalled online via Metrichor’s EPI2ME platform. This generated megabase-scale genomic data for each species (40.4Mbp – 71.7Mbp; see Table 2), with estimated genomic coverage ratios ranging from approximately 3% (*S. uniflora*) to 24% (*N. alata*). N50 sizes of 1.6-2.7 Kbp and maximum read lengths from 66Kbp-144Kbp highlight the exponential distribution of read lengths. Raw reads from each species were compiled into a separate BLAST (NCBI-BLAST+; v2.5.0) nucleotide database.

**Table 2:** R4IDs training-run sequencing statistics. Sequencing runs were each conducted on a separate flowcell, for approximately 24 hours’ duration each.

| Species | Total reads | Total yield | Mean | Median | Max | N25 | N50 | N75 |
| --- | --- | --- | --- | --- | --- | --- | --- | --- |
| *Nepenthes alata* | 51,608 | 68,411,676 | 1,326 | 831 | 95,184 | 3,934 | 2,362 | 1,280 |
| *Sorbus aria* | 36,417 | 51,914,413 | 1,426 | 854 | 66,551 | 4,485 | 2,705 | 1,452 |
| *Beta patula* | 23,694 | 40,492,055 | 1,709 | 1,194 | 117,469 | 4,334 | 2,671 | 1,486 |
| *Silene uniflora* | 24,744 | 40,125,514 | 1,657 | 1,180 | 99,060 | 4,180 | 2,461 | 1,364 |
| *Erycina echinata* | 57,701 | 71,714,205 | 1,243 | 910 | 144,298 | 2,884 | 1,685 | 999 |
| Mean: | <u>38,833</u> | <u>54,531,573</u> | <u>1,472</u> | <u>994</u> | <u>104,512</u> | <u>3,963</u> | <u>2,377</u> | <u>1,316</u> |
| Total: | <u>194,164</u> | <u>272,657,863</u> |  |  |  |  |  |  |

The following week, live at the Kew Science Festival, libraries were prepared from extracted DNA samples and resequenced, two per day. Sample selection was randomised without replacement by an assistant (AJH) such that the sample identity was blinded to the experimenters (JP and ASTP). Live basecalling was used to generate results in real-time. Reads were continuously processed by matching each new read to each of the five R4ID species BLAST databases in turn, retaining single best hits with *e*-value cutoff set at 1.0 and default gap / match parameters and word sizes. Cumulative BLAST alignment lengths (’length‘) and number of identities (’nidents‘) were used to infer the identity of the unknown sample, assuming that the longest cumulative aligned read lengths would correspond to the correct sample. A simple graphical user interface (GUI; Figure 1) was displayed over a local network and projected live to science festival visitors. Code for the ID process and web interface is available at http://github.com/lonelyjoeparker/real-time-phylogenomics.

**Figure 1:**
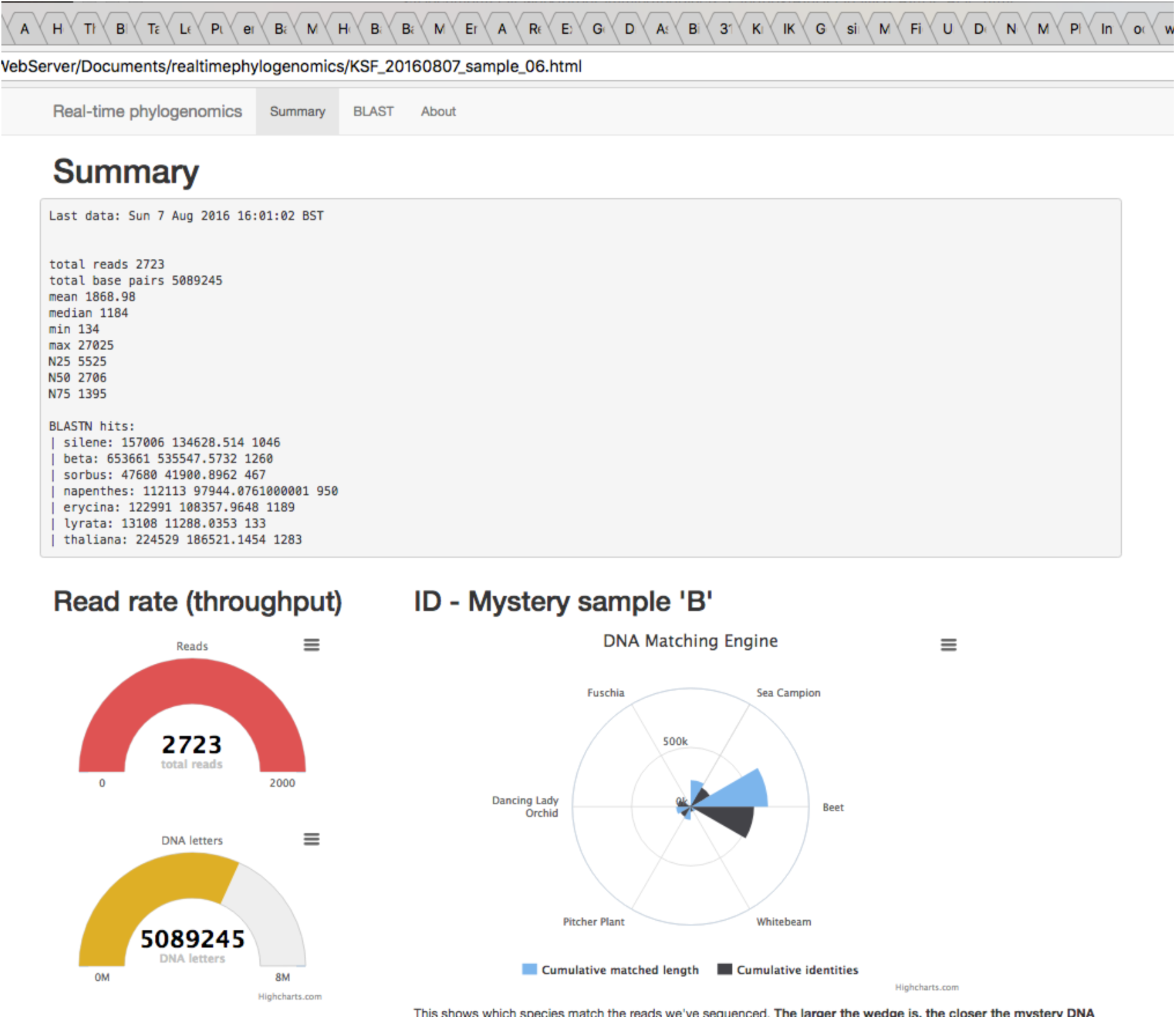
Screenshot of live-updating species ID graphical user interface used at the Kew Science Festival. The text box at the top of the screen displays aggregate BLAST alignment lengths per species, while the graphical plots below display (clockwise from bottom-left) total sequencing yield (bp); number of reads sequences; and a radar segment plot visualising aggregate BLAST alignment length information for total length (light blue segments) and number of aligned identities (charcoal segments). It can be appreciated that BLAST matches are greatly enriched for one species (*B. patula*) over the others – correctly in this case, enabling source sample ID in 15 minutes’ sequencing time.

All five species’ identities were correctly inferred, with the fastest sample load-to-correct guess time being approximately 15 minutes (Table 3). Further resequencing was continued in the laboratory each night to collect further data; total sequencing yields for sample resequencing approximately mirrored those for R4ID training-run sequencing (Table 4), with the exception of *S. aria*, which could not be restarted later owing to time constraints.

**Table 3:** Science festival species-to-blinded-sample mappings, and live-sequencing ID performance (reads sequenced at the point that a correct ID guess was made; for reference approximate R9 MinION throughput with this kit was 5,000 – 10,000 reads per hour.)

| Sample origin | Resequencing run number | Randomised letter | Inferred species ID | Reads sequenced at ID decision | Yield sequenced at ID, bp |
| --- | --- | --- | --- | --- | --- |
| *Nepenthes alata* | Sample_01 | C | *N. alata* | 1,460 | 1,495,967 |
| *Silene uniflora* | Sample_03 | F | *S. uniflora* | 1,421 | 2,725,090 |
| *Erycina echinata* | Sample_04 | E | *E. echinata* | 1,679 | 2,028,975 |
| *Sorbus aria* | Sample_05 | D | *S. aria* | 210 | 505,043 |
| *Beta patula* | Sample_06 | B | *B. patula* | 3,546 | 7,226,106 |
| <u>Mean:</u> |  |  |  | <u>1,193</u> | <u>1,688,769</u> |

**Table 4:**
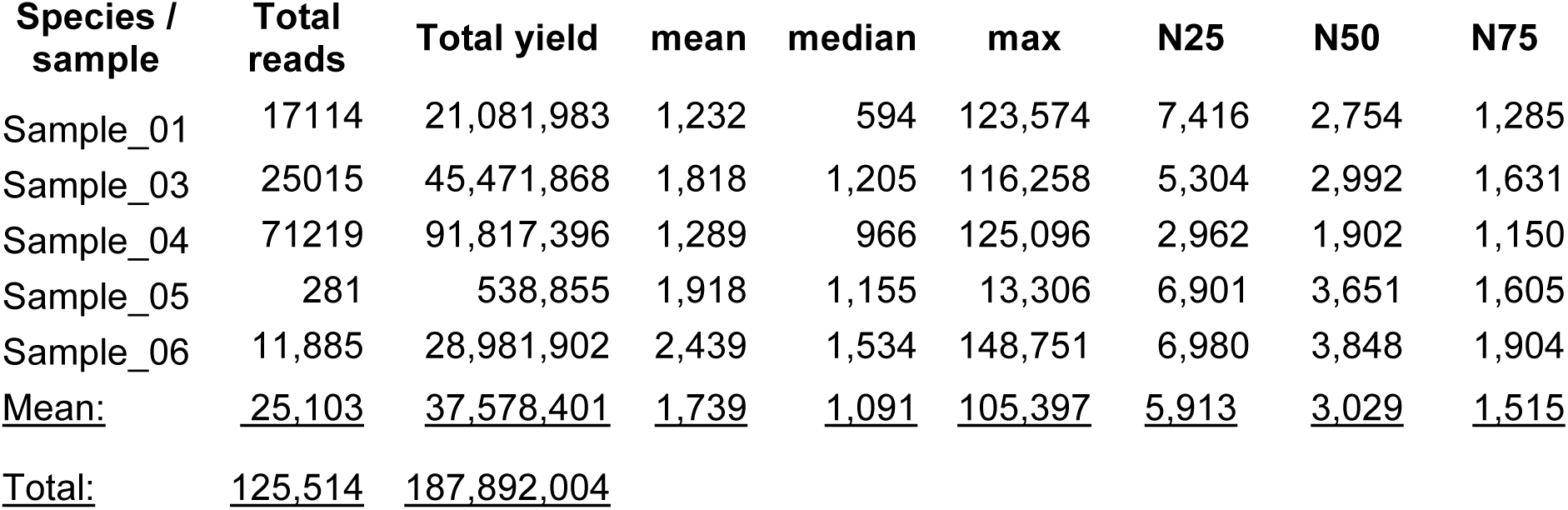
Total resequencing run yields; following a successful live-sequencing ID, flowcells were transported back to the Jodrell Laboratory where resequencing was restarted and completed overnight.

| Species / sample | Total reads | Total yield | mean | median | max | N25 | N50 | N75 |
| --- | --- | --- | --- | --- | --- | --- | --- | --- |
| Sample_01 | 17114 | 21,081,983 | 1,232 | 594 | 123,574 | 7,416 | 2,754 | 1,285 |
| Sample_03 | 25015 | 45,471,868 | 1,818 | 1,205 | 116,258 | 5,304 | 2,992 | 1,631 |
| Sample_04 | 71219 | 91,817,396 | 1,289 | 966 | 125,096 | 2,962 | 1,902 | 1,150 |
| Sample_05 | 281 | 538,855 | 1,918 | 1,155 | 13,306 | 6,901 | 3,651 | 1,605 |
| Sample_06 | 11,885 | 28,981,902 | 2,439 | 1,534 | 148,751 | 6,980 | 3,848 | 1,904 |
| <u>Mean:</u> | <u>25,103</u> | <u>37,578,401</u> | <u>1,739</u> | <u>1,091</u> | <u>105,397</u> | <u>5,913</u> | <u>3,029</u> | <u>1,515</u> |
| <u>Total:</u> | <u>125,514</u> | <u>187,892,004</u> |  |  |  |  |  |  |

### Evaluation of empirical R4IDs

To determine the expected performance of our five-way BLAST ID method at varying levels R4ID training DB and resequencing ‘sampling‘, Python scripts were used to resample the R4ID and resequencing datasets, without replacement, at five levels each: 100, 500, 1000, 5000 and 10,000 reads–producing a total of 25 combinations. This was repeated for each of the five species and 20 replicates, for 625 separate BLAST searches per replicate, collecting alignment length, % identities and *e*-values for each hit. In every case, aggregate hit alignment length (the statistic used in the science festival live-sequencing GUI) favoured the correct species, regardless of R4ID / training run sampling or resequencing effort (see Figures 2 and 3). To analyse the expectation of correct true positive read identifications in more depth, BLAST outputs were parsed to collect the following information: total BLAST hits; one- and two-way true positive hits (reads matching only the correct database, or with longest aligned length to that database); one- and two-way false positive hits (reads matching only an incorrect species database, or with the longest match to an incorrect species); mean length bias (read-wise aligned length for correct species - next longest species aligned length); two-way hit correct percentage (number of two-way hits correctly classified if a length bias > 0 was used for classification); and two-way hit correct % with 50bp threshold (e.g. reads only classified if length bias > 50bp.) Scripts and R Markdown notebooks for these analyses are available online at https://github.com/lonelyjoeparker/oddjects-sandbox/tree/master/R4IDs.

**Figure 2:**
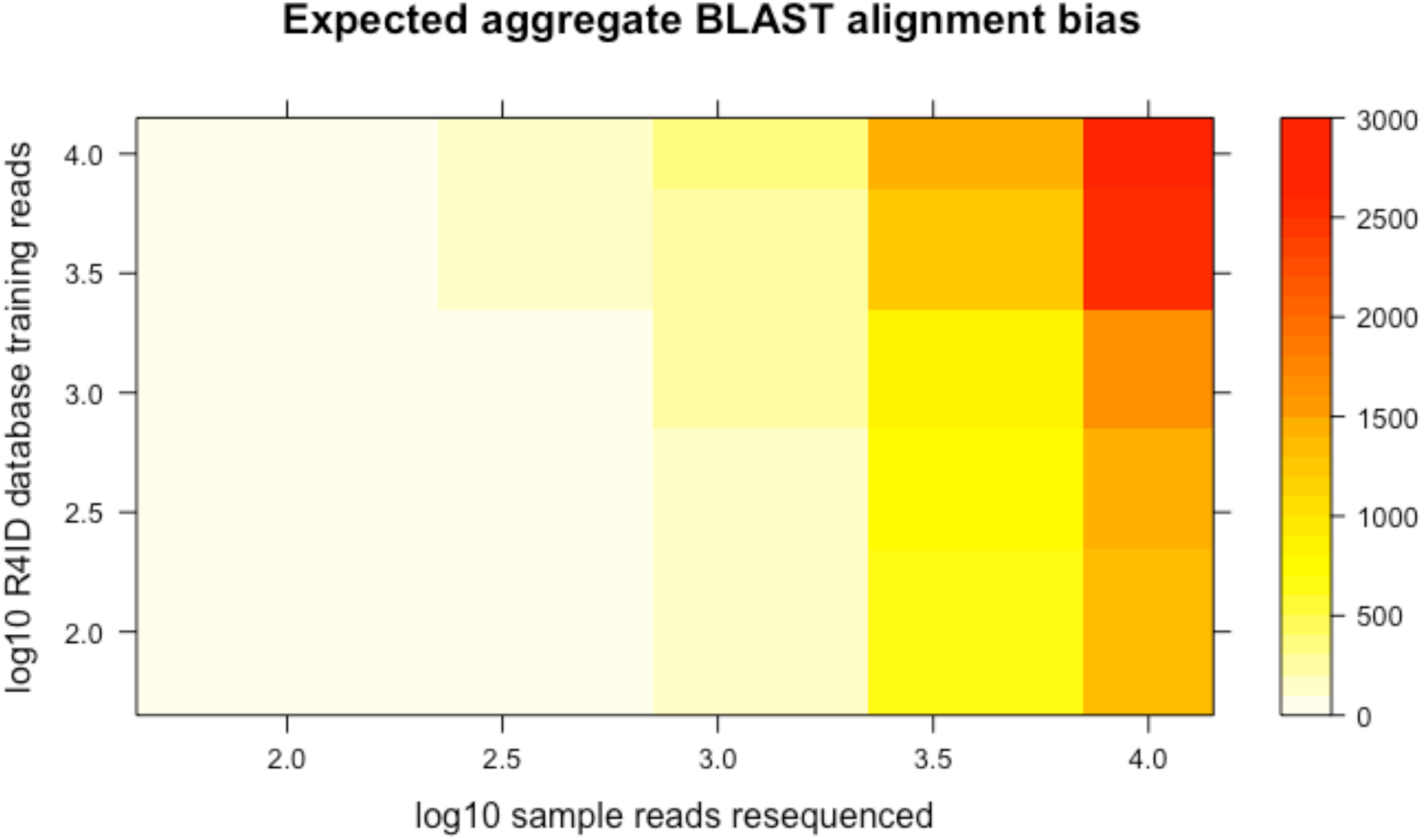
Expected aggregate alignment length bias (total aligned hit length for correct species R4ID database less aligned length for the next closest match) by ‘unknown’ sample resequencing effort (number of reads resequenced; x-axis, log10 scale) and R4ID database training sequencing effort (number of reads sequenced; y-axis, log10 scale). 20 replicates were generated *in silico* by resampling without replacement from the sequenced data.

**Figure 3:**
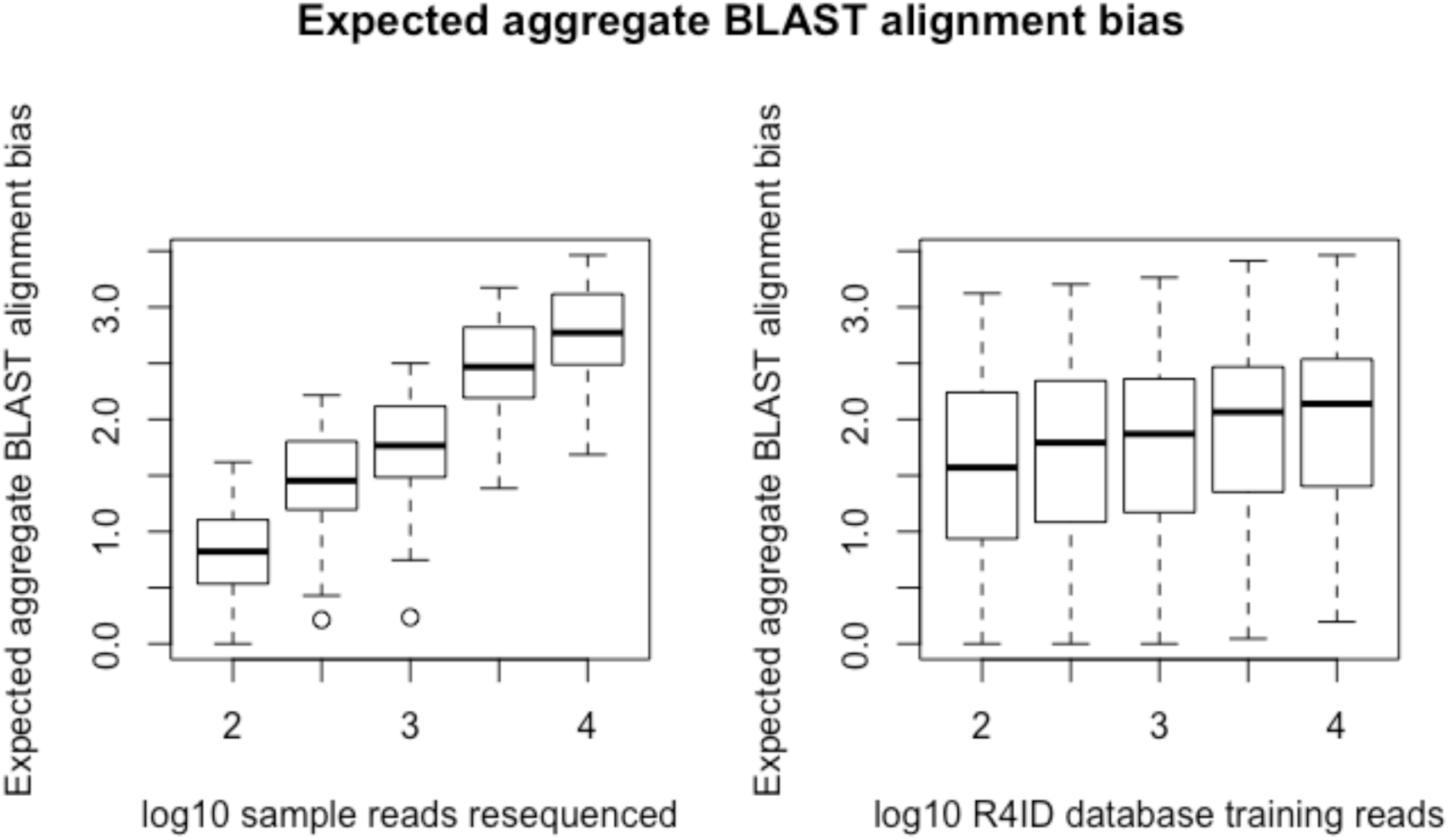
Boxplots showing expected aggregate BLAST alignment length bias (total aligned length for ‘correct’ species R4IDs database vs. next-longest match; y-axis, log10 scale) versus sample resequencing effort (# reads: left plot; x-axis, log10 scale) or R4ID training sequenced effort (# reads: right plot; x-axis, log10 scale).

In order to assess the capacity for individual reads to provide accurate species ID, we calculated the per read accuracy (ratio of true positives to true & false positives; Figure S1). This ratio exceeds 50% at most sampling levels, including as low as 100 reads, however, the pattern is principally driven by R4ID training effort, with 5,000 or more reads likely to be sufficient for >60% correct IDs per read. We also estimated the expected total number of positive reads as a function of total resequencing effort when the threshold for the aligned length bias statistic used for species assignment is set to greater than zero bp (less conservative; Figure S2a) or greater than 50bp (more conservative; Figure 5b).

**Figure 4:**
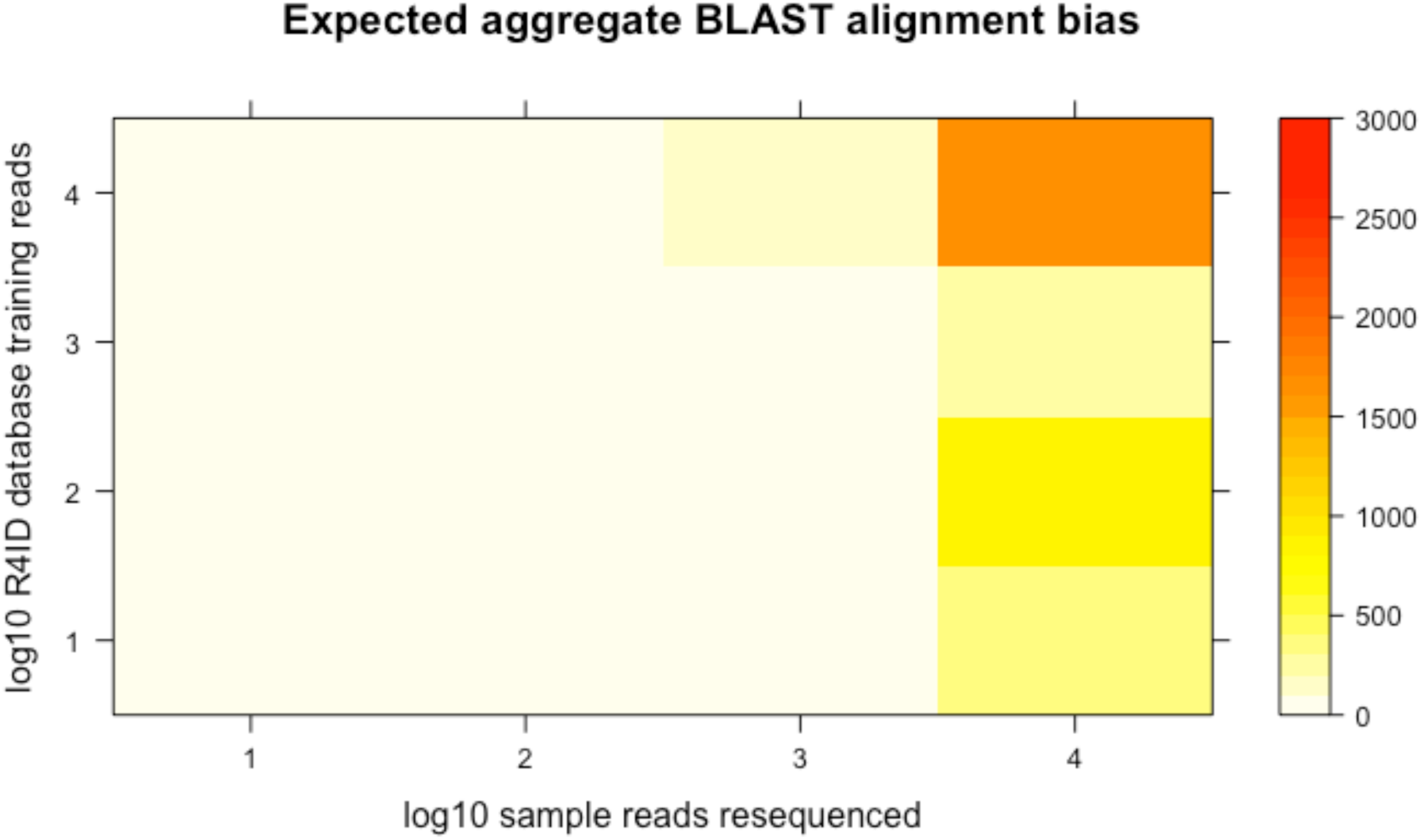
Expected aggregate alignment length bias (total aligned hit length for correct species R4ID database less aligned length by ‘unknown’ sample resequencing effort (number of reads resequenced; x-axis, log10 scale) and R4ID database training sequencing effort (number of reads sequenced; y-axis, log10 scale). 18 replicates were generated *in silico* by sampling published genome assemblies using an error model derived from empirical nanopore sequencing data (see text for details).

**Figure 5:**
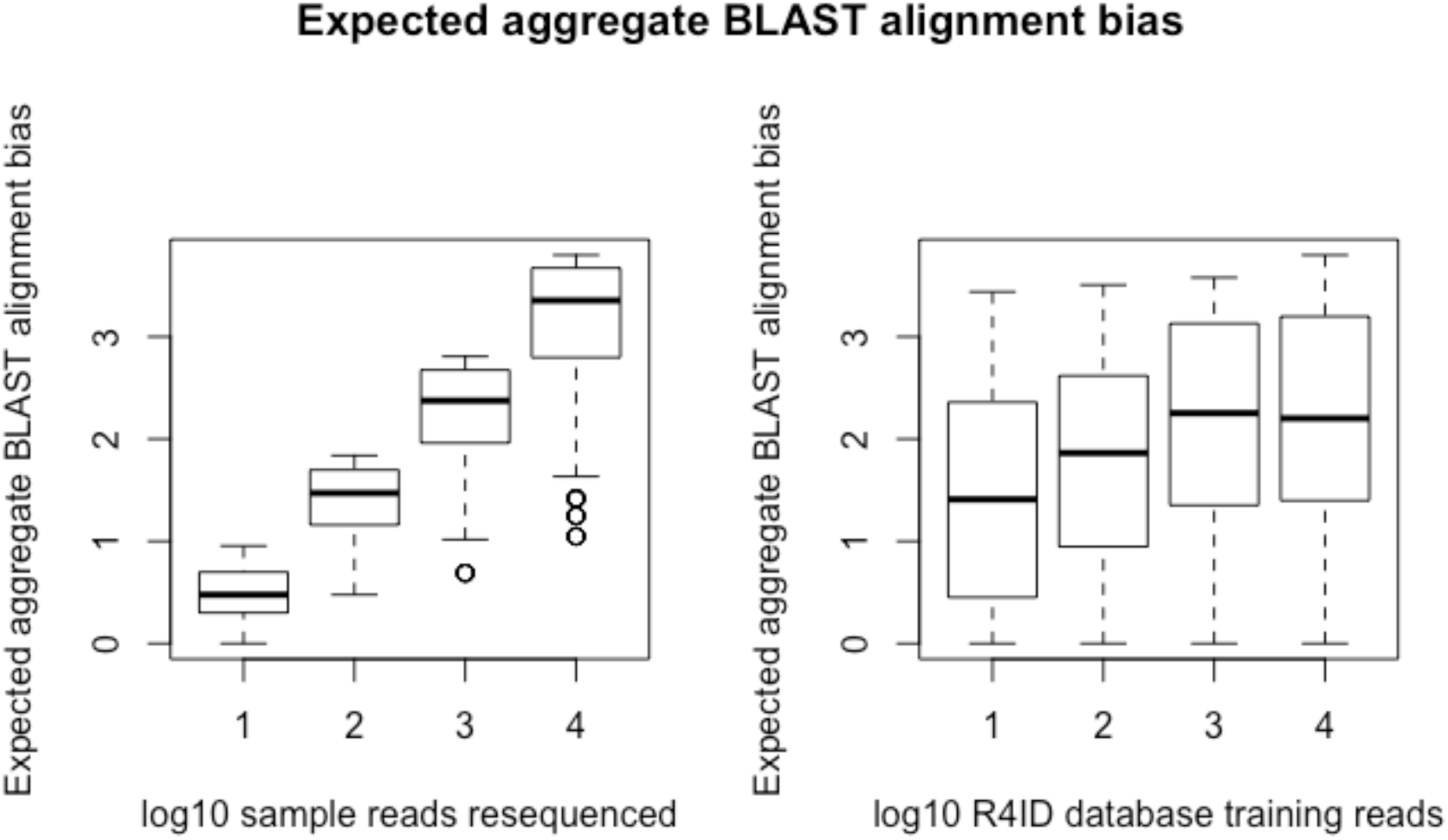
Boxplots showing expected aggregate BLAST alignment length bias (total aligned length for ‘correct’ species R4IDs database vs. next-longest match; y-axis, log10 scale) versus sample resequencing effort (# reads: left plot; x-axis, log10 scale) or R4ID training sequenced effort (# reads: right plot; x-axis, log10 scale).

### Evaluation of simulated R4IDs

The five species sequenced were distributed widely across the vascular plant phylogenetic tree. To evaluate likely performance of our method on a more closely related group of species, we performed simulated nanopore sequencing of six species from the Cammelinae with publically available genomes (details given in Table 5); *Arabidopsis thaliana, A. lyrata* and *A. halleri, Capsella rubella, C. bursa-pastoris* and *Camelina sativa.* Assembled genome lengths for these species range from ∼ 120Mbp (7 complete chromosomes;*A. thaliana*) to ∼640Mbp (37,780 contigs; *C. sativa*). To simulate MinION 1D rapid sequencing runs (for both R4ID training sequencing or sample ID resequencing), 10, 100, 1000 or 10,000 reads were drawn randomly from the genome assemblies. Contigs and starting positions were selected uniformly, and read lengths such that they were exponentially distributed with an N50 approximately equal to 1kbp).

**Table 5:** Statistics of public genome sequences used in *in silico* R4IDs simulation. *Concatenated for this analysis.

| Species | Accession | Assembly length, Mbp | Number of contigs |
| --- | --- | --- | --- |
| *Arabidopsis thaliana* | TAIR10 | 119 | 7 |
| *Capsella rubella* | ANNY01.1 | 129 | 7,067 |
| *Arabidopsis halleri*<br>*subsp. gemmifera* | FJVB01.1 | 196 | 2,239 |
| *Arabidopsis lyrata* | ADBK01000001.1 | 207 | 695 |
| *Capsella bursa-pastoris* | MPGU01.1 | 268 | 8,186 |
| *Camelina sativa* | JFZQ01.1;<br>JFZQ01.2* | 641 | 37,780 |

The MinION sequencing platform is known to have a higher error rate than lab-based next-generation sequencing. To approximate this, we first analysed the error characteristics of alignable *A. thaliana* reads from our previous experiments (Parker *et al.* 2017) and derived a simple error model. This assumes an error rate of 5% such that one in 20 positions were selected randomly with replacement. Errors were randomly assigned as either substitutions (direct point substitution being drawn from all four bases equally), deletions, or homopolymer insertions (the *n+1th* base being inserted as a copy of the *n*th base), since homopolymer errors are a comparatively common occurrence on this platform. Having repeated this process for 18 replicates, a total of 576 total combinations (4 resequencing sample levels * 4 R4ID levels * [6 * 6 species comparisons]) was analysed with BLASTN as above. Code is available for the nanopore sequencing simulation at the above Github repository.

The outcome of the simulated R4ID experiments are displayed in the same format as for the empirical experiment (Figures 4 and 5). As with the empirical and resampled data from wider phylogenetic comparisons, this group of closely-related species showed the expected aggregate length biases that were greater than zero even for comparatively low amounts of R4ID sequencing and sample resequencing effort (1000 or more reads in each case).

Again, the R4IDs database sequencing effort was a principal driver of accuracy. Figure S3 shows how true- and false-positive hit rates vary with number of reads sequenced for the R4ID database; once 10,000 reads have been sequenced, the true-positive hit rate (∼0.55) considerably exceeds the false-positive rate (<0.2).

Since the assembled genome lengths for these closely-related species vary from ∼100Mbp to ∼650Mbp, we examined how this affected the ID rate for each species (Figure 9). A greater number of hits were observed at lower sequencing sampling effort for *A. thaliana* (the smallest genome, at approximately 119Mbp) compared with *C. sativa* (the largest, at 641Mbp). This is unsurprising, since each R4ID database of a given size represents a lower fraction of the genome as the genome size increases.

Our real-time analysis pipeline has been packaged up as deployable Docker containers, making it simple for anyone to generate R4IDs (’training‘), evaluate resequenced data (’resequencing‘) and display results on a local web server (’visualisation‘). These are available from Docker Hub at https://hub.docker.com/r/lonelyjoeparker/r4ids/

## Discussion

We have shown that even partial genomic sequencing using long-read nanopore data provides an adequate training dataset for future resequencing. This demonstrated that numerous potential uses for DNA sequence data that are impeded by availability of reference sequences may now be exploited with the rapid generation of non-specific genomic sequencing. Intuitively, this may seem unsurprising, but our results clearly demonstrate how little data is needed for a successful sample ID. The sequencing effort is low both in terms of sample resequencing and more importantly in terms of the ‘reference’ genome sketch generated from a sample of known origin. In each case, our empirical, resampling, and simulation results (including closely-related species) show that as few as 10,000 reads are easily sufficient to achieve reliable sample ID with nanopore long-read sequencing.

At the time that the empirical data was collected (August 2016), 30-120 minutes’ sequencing were required per species/sample. MinION platform yields have greatly improved since then, as has the data accuracy - Oxford Nanopore’s informal ‘Poreboard’ competition currently stands at >11Gbp sequencing from a single flowcell, while the longest single mapped read now stands at 1.3Mbp with 97% accuracy (M. Loose, pers. comm.). In this context, training datasets for multispecies R4ID matching could easily be collected in a few hours’ sequencing. To identify unknown samples by resequencing and comparison against previously-sequenced R4ID databases even more rapid. Our live sequencing at the Kew Science Festival 2016 managed 5 out of 5 correct identifications of blinded samples in as little as 15 minutes. Recent improvements in platform throughput suggest that this time is likely to fall in the near future.

We observed three main drivers of identification efficiency and accuracy. First, the level sequencing effort employed in the generation of the R4IDs training datasets. Second, the size of the target genome, since this will interact with sequencing effort to give the expected genome coverage fraction. Thirdly, the phylogenetic distance between candidate species: our expected hit fractions were lower, though still acceptable, in our *in silico* simulations on closely-related species. Our overall approach is well-validated by these empirical and in silico experiments. One possible reason for the success of this method may be the abundance of plastid and other organelle material present in extracted DNA. Outline genomic mapping analyses using this data and data from our previous work (Parker *et al*, 2017) show a high prevalence of chloroplast genomic material with nearly complete plastids being assembled post hoc. This is unlikely to be the sole cause as whole plastid genome sequencing is demonstrably not a panacea for species identification, however, the combination of abundant plastid sequences and more limited matching to nuclear data may be an important factor. In addition to the phylogenetic distance mentioned larger phylogenetic distances in the empirical dataset, this detail may explains why the empirical data outperform the simulated data in key respects.

Typical applications for this approach could include rapid digitisation of collections and biodiversity evaluations, particularly where applications and sampling opportunities are limited to on-site data collection (perhaps for CITES permit reasons) or time-limited (as in the case of transient and unexpected events, such as the *Deepwater Horizon* rig fire and subsequent contamination of the Gulf of Mexico (partially sampled *post-hoc* by Mason *et al.*, 2014) Our method is inherently real-time and, with some extension, might allow in-the-field species delimitation or phylogenomic inference. Finally, we note that the computational requirements of this technique are actually comparatively light, and could even be met by a small array of embedded single-board machines such as the Raspberry Pi (Barker *et al.*, 2013) or Parallela devices. This in turn enables field-sequencing at remote locations where data uplink may be limited, such as hydrothermal vents (Smedile *et al.*, 2013; Dick & Tebo, 2010), deep-sea brine pools (Wang *et al.*, 2013), or pelagic sampling approaches which have previously been limited by the availability of shipboard facilities for sequencing or sample storage (Ventner *et al.*, 2004).

In contrast to molecular barcoding and hybrid-capture approaches, our method does not require existing primers or particularly complicated labwork, and is inherently portable for field-work and scalable to completely novel groups. Furthermore, the data from other barcoding approaches can easily be incorporated into R4ID databases, building upon the valuable and highly curated barcode datasets that are already available. Molecular systematics based on multiple loci also have clear advantages over barcode-based approaches, since they allow models which account for reticulate evolution and horizontal gene transfer (Pryron, 2015; Mallo & Posada, 2016). In addition, it has the clear advantage that truly genomic data can be captured both in the training and resequencing phases, for future use. This is important, because it can reasonably be expected that low-coverage data sequenced for R4ID training or sample identification can yield useful downstream data for genomic, metagenomic or phylogenomic studies With improving read accuracy collecting these kinds of broad genomic sketches/skims could revolutionise the way in which we delimit species and gather population genetic data.

## Acknowledgements

This work was carried out at the Kew Science Festival 2016, with support from the Kew Foundation. Oxford Nanopore provided flowcells, reagents and technical support free-of-charge but have had no editorial input into downstream analyses, or preparation of this manuscript. We are grateful to Kew colleagues in the Science Directorate for assistance with sequencing logistics, and in the Horticultural Directorate for assistance with sample collection; and to Gaetan Lee and colleagues for assistance and funding for the Kew Science Festival exhibit.

